# Osmotic stress triggers fast and reversible PMF collapse in *Escherichia coli*

**DOI:** 10.1101/2025.09.25.678644

**Authors:** Luis Meneses, Eric M. Dudebout, Sophia Belser, Jinming Yang, Navish Wadhwa

## Abstract

Environmental stressors routinely impact bacteria and affect their physiology. Among the most important physiological parameters is the proton motive force (PMF), that powers vital cellular processes and molecular machines. Measuring how PMF responds to environmental stress in real time requires tools that capture rapid physiological changes with high temporal resolution. Here, we use the bacterial flagella motor as an *in vivo* voltmeter to probe PMF dynamics during hyperosmotic shock. Because motor rotation frequency scales linearly with PMF, this approach enables single-cell electrophysiology with high temporal resolution. We find that hyperosmotic stress causes a rapid and reversible loss of PMF in *Escherichia coli*, independent of the osmolyte used or the presence of potassium ions. We corroborate these findings by using the Nernstian dye, tetramethyl rhodamine methyl ester (TMRM), showing that hyperosmotic shock leads to membrane depolarization. Together, our results highlight the efficacy of the flagellar motor as powerful tool for probing bacterial electrophysiology and reveal that hyperosmotic stress directly disrupts cellular energetics in addition to its mechanical effects.

## INTRODUCTION

Bacteria encounter a wide range of environmental stimuli and must exhibit tailored adaptive responses to maintain homeostasis and promote survival (Dufrêne and Persat, 2020). In addition to chemical stimuli, bacterial cells routinely experience mechanical stressors that affect their physiology (Belas, 2014; Persat, 2017). Although bacterial responses to chemical cues are well characterized, uncovering how cells respond to mechanical stress in real time represents a critical next step in improving our understanding of microbial behavior (Gordon and Wang, 2019).

PMF is a key biological parameter in bacteria, derived from an electro-chemical gradient of protons across the inner membrane. It powers vital cellular processes, including cell division (Strahl and Hamoen, 2010), biofilm formation (Prindle et al., 2015), ATP synthesis (Mitchell, 1961; Maloney et al., 1974) and the flagellar motility (Manson et al., 1977; Wadhwa and Berg, 2022). Because of its essential role, the PMF is generally believed to be homeostatic, though its steady state value may depend on physio-chemical factors such as nutritional status and metabolic demand (Benarroch and Asally, 2020). However, recent studies showed that PMF is highly dynamic on short time scales (Kralj et al., 2011; Biquet-Bisquert et al., 2024). In the absence of stimuli, PMF can undergo transient rapid fluctuations (Kralj et al., 2011). Chemical signals, such as potassium fluctuations from intercellular communication in biofilms (Prindle et al., 2015; Akabuogu et al., 2025), and physical stimuli, such as surface contact (Bruni et al., 2017), can also lead to dynamic changes in PMF. However, despite growing evidence of PMF’s sensitivity to internal and external cues, its real-time behavior under environmental stress has yet to be fully elucidated.

Measuring bacterial PMF presents a difficult challenge. Patch clamp techniques, used for electrophysiological measurements in eukaryotic cells, cannot be applied to bacteria due to their small size and the presence of a cell wall (Benarroch and Asally, 2020). Instead, optical, fluorescence-based methods employing dyes and fluorophores are commonly used (Mancini et al., 2020). These include Nernstian dyes, that label the cell in a membrane potential-dependent manner, and genetically encoded voltage indicators, whose fluorescence emission responds to local membrane potential (Jin et al., 2023). Although widely used, these optical methods have distinct limitations, such as the need for careful calibration protocols, ratiometric reporters, and, in case of Nernstian dyes, permeabilization of the outer membranes of Gram-negative bacteria (Krasnopeeva et al., 2019; Buttress et al., 2022). These limitations necessitate the development of alternative approaches to measure PMF in bacteria.

The bacteria flagellar motor can be used as a biological voltmeter to measure real-time changes in the PMF without the limitations of fluorescence-based approaches (Krasnopeeva et al., 2019; Biquet-Bisquert et al., 2021, Biquet-Bisquert et al., 2024). Rotation of the flagellar motor is powered by proton translocation across stators embedded in the inner membrane (Wadhwa and Berg, 2022). The rotation speed of the motor is directly proportional to the PMF, allowing one-to-one measurement between the two (Gabel and Berg, 2003). Therefore, measuring the rotational speed of the flagellar motor provides high temporal resolution measurement of bacterial electrophysiology *in vivo*.

Among the environmental stressors experienced by bacteria, osmotic stress is one of the most ubiquitous. A sudden change in external solute concentration drives rapid water flux across the inner membrane, leading to significant mechanical deformation of the cell (Pilizota and Shaevitz, 2013; Fuchino et al., 2017). This deformation can lead to drastic outcomes, including plasmolysis (separation of the inner membrane from the cell wall) (Cota-Robles, 1963) and even cell lysis (Wong and Amir, 2019). Cellular responses to osmotic stress have been extensively studied (Sleator and Hill, 2002), and many of the underlying pathways are now well characterized (Altendorf et al., 2009). However, experiments involving osmotic stress are typically conducted with time resolutions of minutes to hours and often involve growing bacteria at different osmolarities. In the environment, osmolarity can shift dramatically within seconds, such as during flooding or surface runoff that dilutes bacterial microenvironments. While prior research has illuminated how cells cope with osmotic stress over time, their immediate response following a sudden shock is still not fully understood.

Here, we use the flagellar motor as a voltmeter to investigate how the PMF is affected by a hyperosmotic shock. We show that a hyperosmotic shock rapidly diminishes motor rotation speed in a manner that depends on the shock strength. The rotation loss is independent of the choice of osmolyte, indicating that the observed response is broadly applicable to diverse osmotic stimuli. The decrease in rotation occurs even in the absence of potassium, precluding potassium uptake as the underlying mechanism. The response upon hyperosmotic stress also occurs in clockwise-locked motors, indicating that the loss of motor rotation does not depend on specifics of the stator-rotor interaction within the flagellar motor. Finally, we show that the loss of motor rotation is associated with a loss of membrane potential by corroborating our results with a Nernstian fluorescent dye, tetramethyl rhodamine methyl ester (TMRM) (Lo et al., 2007; Mancini et al., 2020). In sum, we discuss the use of the flagellar motor as a sensor for monitoring a cell’s physiological state and expand the developing field of bacterial electrophysiology by demonstrating that *E. coli* cells undergo depolarization in response to hyperosmotic stress.

## RESULTS

### Hyperosmotic shock causes rapid reduction of motor rotation speed

Our experiment is based on the flagellar motor bead assay, in which a micron-sized bead is attached to the flagellar filament and its motion is recorded as a readout of motor rotation. However, instead of probing motor dynamics, we use bead assays to investigate the PMF. The rotation speed of the flagellar motor is proportional to the PMF; motor rotation occurs with proton flux across stator units (Fig. 1A). Therefore, motor rotation speed is a direct measure of the PMF, making the flagellar motor an *in vivo* voltmeter.

**Figure 1.**
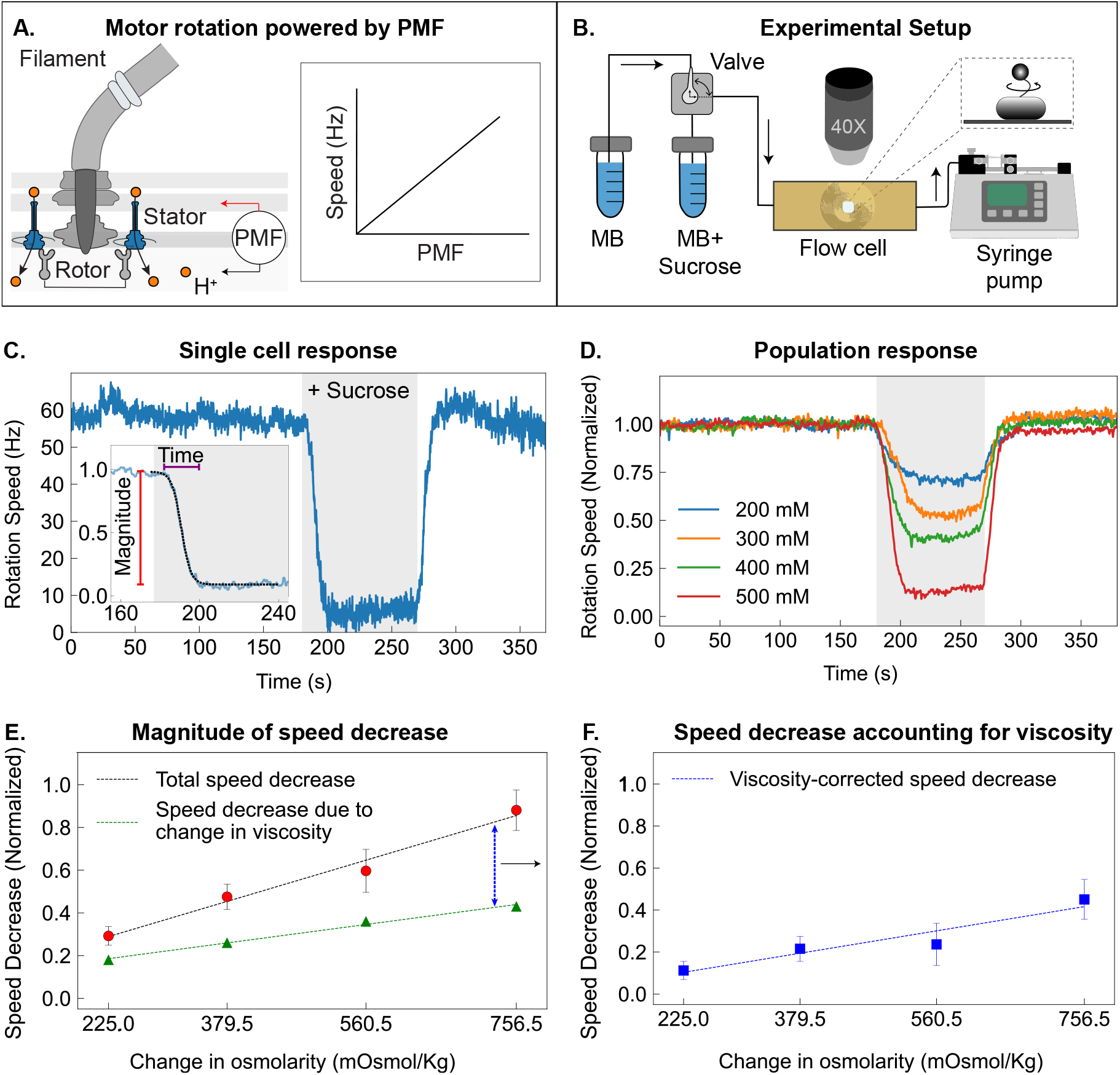
Bacterial flagellar motor response to hyperosmotic shock measured by bead assay. **A)** Schematic of the motor, where proton motive force (PMF) drives rotation via stator units; motor speed is linearly proportional to PMF (inset). **B)** Bead assay: A fluidics system alternates between motility buffer (MB) and MB + sucrose. A 1 μm bead attached to the *E. coli* filament tracks motor rotation, imaged with a 40X phase contrast microscope and a high-speed CMOS camera. **C)** Example trace showing motor speed drop during 500 mM sucrose shock (gray) and recovery after removal. Inset: normalized trace with a sigmoidal fit showing the magnitude (red bar) and the timescale (purple bar) of speed reduction. **D)** Population-averaged responses to 200-500 mM sucrose shocks. Larger shocks result in greater speed reductions. *n* = 8, 10, 8, and 12, for 200 mM, 300 mM, 400 mM, and 500 mM shocks respectively. **E)** Normalized reduction in motor speed vs. change in osmolarity. Red circles and black bars represent the mean and standard deviation (SD), respectively. The dashed black line is a linear fit. Green triangles and the dashed green line represent the estimated speed decrease due to the increase in viscosity and a linear fit to those data. The blue dashed double arrow represents the difference between the total and viscosity-driven speed decreases, plotted in the next panel. **F)** Viscosity-corrected speed decreases. Blue squares and error bars represent mean and SD, respectively. The dashed blue line is a linear fit. The parameters for the fits are provided in Table S3 and Table S4.

To probe the PMF response to a hyperosmotic shock, we immobilized *E. coli* cells inside a custom-made flow cell (Berg and Block, 1984) and attached 1 μm polystyrene beads to their flagellar stubs. We used an *E. coli* strain expressing a mutant filament protein (FliC-st) to which beads readily adhere. This strain also lacked the chemotaxis response regulator CheY and therefore rotated its motors exclusively counterclockwise. Under a constant flow rate set by a syringe pump, the media entering the flow cell could be switched from isotonic to hyperosmotic using a manual valve. The cells were initially kept under a constant flow of motility buffer (MB). When MB was replaced with hyperosmotic MB + sucrose, the immobilized bacteria inside the flow cell experienced a temporary osmotic shock (Fig. 1B).

An osmotic shock resulted in a rapid loss in the rotation speed of the flagellar motor. We began the experiment by recording the initial rotation speed of an individual motor in MB (219 mOsmol/Kg) for 180 s. We then applied a hyperosmotic shock, indicated by the shaded region in (Fig. 1C), by exchanging the media with MB supplemented with 500 mM sucrose. Prior to the shock, the motor rotated at a stable speed of approximately 50 Hz. Immediately after the shock was applied, motor speed decreased to approximately 10 Hz. During the shock, motor rotation did not cease and remained approximately constant at its minimum (Fig. 1C). At 270 s, we replaced the MB + sucrose media with MB, removing the hyperosmotic shock. Motor speed quickly recovered to its original value and remained constant for the duration of the recording.

Across different osmotic shock strengths, single motor speeds varied in magnitude but followed similar dynamics (Fig. S1). We tested the cell’s response to MB supplemented with 200 mM, 300 mM, 400 mM, and 500 mM sucrose (Fig. 1D), corresponding to changes in osmolarity of 225, 379.5, 560.5, and 756.5 mOsmol/Kg, respectively (Table S5). For each condition, we normalized the motor speeds to their pre-shock values and averaged across multiple cells. Higher sucrose concentrations caused greater reductions in the population-averaged motor speed (Fig. 1D). We quantified the magnitude and the timescale of motor speed decrease by fitting an inverse sigmoidal curve to the data (see Methods), showing, as expected, that the magnitude of speed decrease increased with the strength of osmotic upshift (Fig. 1E). In contrast, the characteristic time of the loss in motor rotation speed did not vary significantly with the strength of osmotic shock (Fig. S4).

We ruled out a change in viscosity as the sole cause for the observed change in motor speed. An increase in sucrose concentration of the medium leads to an increase in viscosity (Table S1), which would result in a reduction in the speed of motor rotation. However, the resulting change in viscous drag experienced by the bead was insufficient to explain the observed loss in rotation speed during the hyperosmotic shock (Fig. 1E and Fig. S2). This suggests that hyperosmotic shock induces a second biophysical effect that alters motor speed. We calculated the portion of the speed decrease unaccounted for by viscosity, and found that this difference also increases linearly with the change in osmolarity (Fig. 1F). Thus, while viscosity contributes to the mechanical load on the motor, it does not fully account for the magnitude of speed loss observed during hyperosmotic shock.

Another possible explanation for changes in motor speed is a change in the number of stator complexes driving the motor. Indeed, stators are mechanosensitive and undergo load-dependent remodeling (Lele et al., 2013; Nord et al., 2017; Wadhwa et al., 2019, 2021). However, we did not observe stepwise changes in rotation speed that are a hallmark of stator remodeling (see, e.g., inset in Fig. 1C). Additionally, the observed recovery in rotation speed is significantly faster than that measured for stator remodeling in the same strain (Wadhwa et al., 2019). Thus, we conclude that the change in rotation speed of the motor is likely driven by a loss of PMF in the cell.

### Response to osmotic shock is independent of osmolyte identity

Next, we tested if that the observed response of the flagellar motor to an osmotic shock did not depended on the choice of osmolyte. To do this, we replaced sucrose in our hyperosmotic media with sorbitol and again measured the motor’s response to 200 mM, 300 mM, 400 mM, and 500 mM hyperosmotic shocks. Like sucrose, sorbitol is a non-metabolized, uncharged molecule that enters the *E. coli* periplasm rapidly due to its small size but is taken up into the cytoplasm relatively slowly (Stephens et al., 2024; Rojas et al., 2014). Population-averaged traces of motor speed (Fig. 2A) showed that the response to sorbitol was very similar to sucrose. During the osmotic upshift, motor speed dropped sharply and stabilized at its minimum. After returning to the isotonic MB, speeds quickly recovered to pre-shock levels. Like for sucrose shocks, the characteristic time of motor speed loss was largely independent of the sorbitol concentrations tested, on the order of 5 seconds (Fig. S5A). The similar response to sorbitol and sucrose shocks suggests that motor speed loss during hyperosmotic shock is a general physical phenomenon, not a specific cellular response to particular osmolyte.

**Figure 2.**
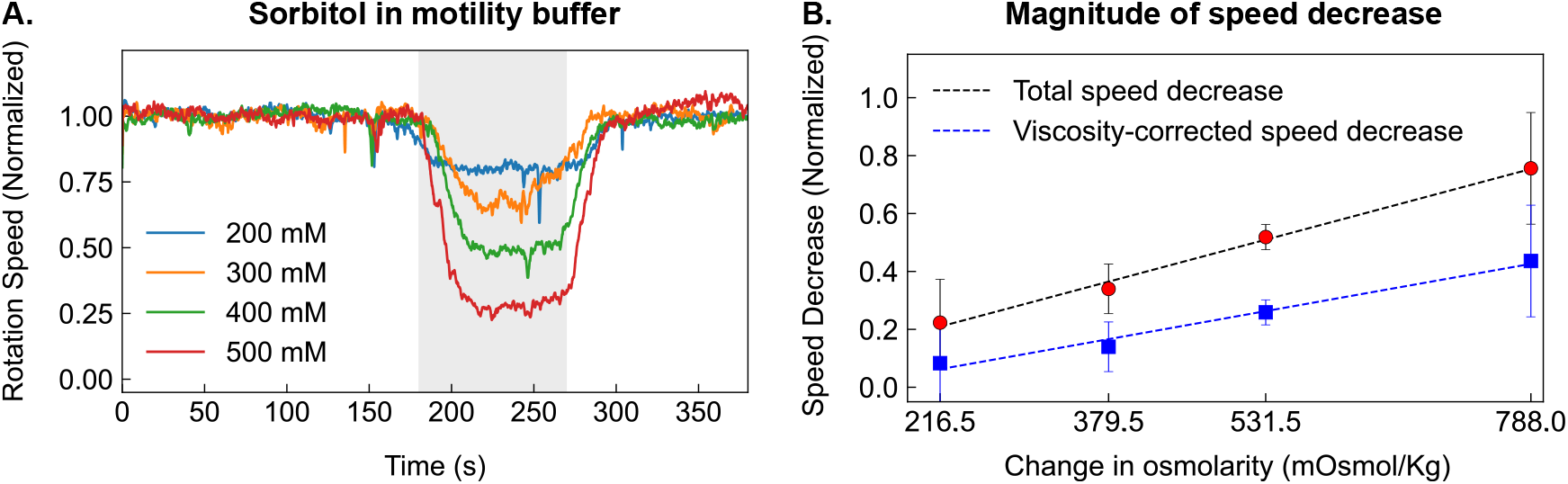
Motor response to osmotic shock is independent of osmolyte identity. **A)** Population-averaged rotation speeds for 200–500 mM sorbitol shocks (*n* = 8 per condition). Shaded region indicates shock duration. **B)** Normalized speed reduction vs. change in osmolarity (from MB with sorbitol). Red circles and black error bars represent the mean and SD, respectively. The dashed black line is a linear fit to the total speed decrease. Blue squares and error bars show the difference between the total speed decrease and the estimated reduction due to changes in viscosity. The dashed blue line represents a linear fit to this difference. Fit parameters are provided in Table S4.

As with sucrose, increasing sorbitol concentration raised the medium’s viscosity, which in turn reduced motor speed (Table S2). Once again, the estimated reduction in rotation speed due to increased viscosity did not fully account for the observed speed loss (Fig. 2B and Fig. S3). Thus, osmotic stress reduces motor speed, likely through dissipation of the PMF. Although the total speed decrease was smaller with sorbitol than with sucrose, correcting for viscosity revealed that the unaccounted for speed decrease was identical for the two osmolytes (Fig. S6). Thus, despite differing effects of viscosity, the corrected speed loss in both conditions is likely due to a common cellular response involving PMF disruption.

### Rotation speed decrease does not require potassium ions

Next, we investigated the role of potassium, a key player in bacterial response to osmotic shock (Stautz et al., 2021). In *E. coli*, response to hyperosmotic shock involves uptake of potassium ions, that counterbalances the loss of turgor pressure, increasing it to homeostatic levels and enhancing survival (Dinnbier et al., 1988). In principle, a rapid influx of positively charged ions could, as a by-product, influence the PMF. We therefore investigated whether a rapid uptake of potassium causes the observed motor response. To test this, we repeated the sucrose osmotic shock experiments with sodium phosphate buffer (SPB), a medium that lacks potassium, as a base instead of MB. Our results showed that the observed motor response to sucrose osmotic shock in SPB closely resembles the motor response to sucrose in MB, indicating that the motor speed loss does not depend on the presence of potassium in the medium (Fig. 3A). As before, accounting for increased viscosity did not fully explain the observed speed loss under SPB sucrose osmotic shock (Fig. 3B). Additionally, the timescale of speed decrease did not depend on the strength of the osmotic shock (Fig. S5B). Thus, while long-term adaptation to hyperosmotic stimuli is aided by the uptake of potassium ions (Dinnbier et al., 1988), the immediate loss in motor speed under hyperosmotic shock still occurs in a potassium-deficient medium, indicating that this response does not involve a rapid potassium influx.

**Figure 3.**
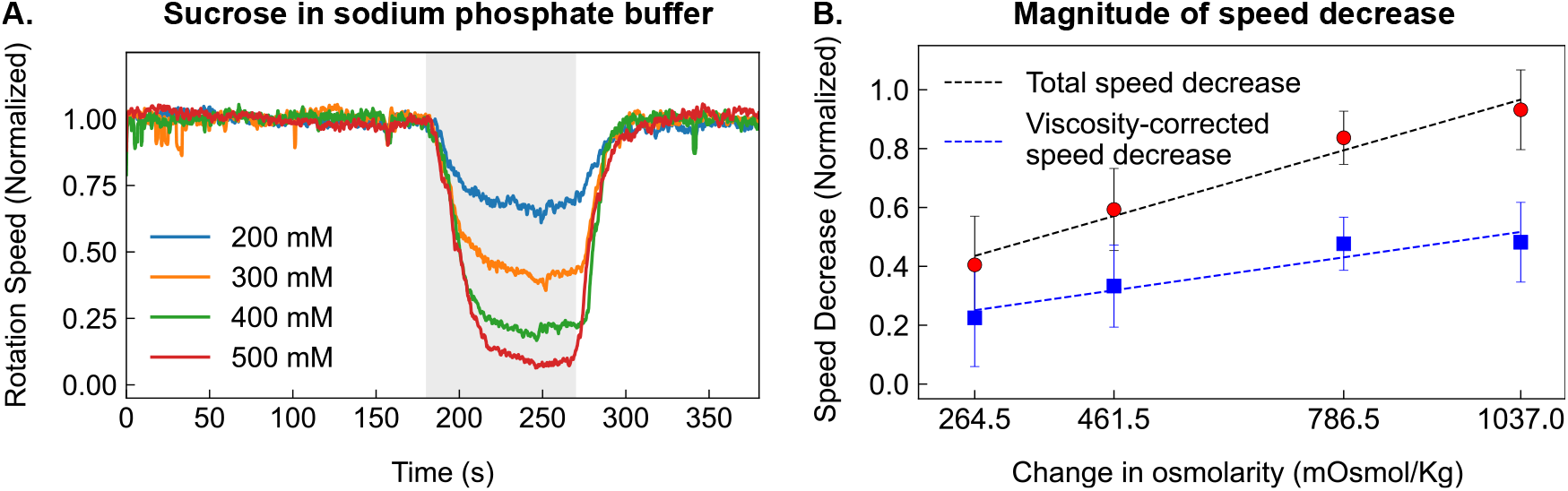
Motor response to osmotic shock is independent of medium. **A)** Population-averaged rotation speeds for 200–500 mM sucrose shocks in sodium phosphate buffer (SPB), with *n* = 10 per condition. **B)** Normalized speed reduction versus change in osmolarity (from SPB with sucrose). Red circles with black error bars indicate the mean and SD, respectively. The dashed black line is a linear fit to the total speed decrease. Blue squares and error bars show the difference between the total speed decrease and the estimated reduction due to changes in viscosity. The dashed blue line represents a linear fit to this difference. Fit parameters are provided in Table S4.

### Motor speed decrease is independent of the direction of motor rotation

A key trait of the bacterial flagellar motor is the capacity to switch between counterclockwise or clockwise rotation. The two configurations have different torque-speed characteristics. In counterclockwise rotation, motor torque decreases slowly with increase in rotation speed, however, once the motor reaches a speed over 150 Hz, the torque decreases sharply (Yuan et al., 2010). In contrast, clockwise rotating motors exhibit a linear decrease of the torque with speed throughout their operational range (Yuan et al., 2010). To determine whether the observed response to osmotic shock depended on motor configuration, we repeated the sucrose osmotic shock experiments with an *E. coli* strain that overexpresses the response regulator

CheY through an IPTG inducible plasmid, and therefore has its motors rotate exclusively clockwise. The observed response closely followed that of counterclockwise rotating motors (Fig. 4A). Once again, under clockwise rotation, the correcting for increased viscosity did not fully explain the observed speed loss (Fig. 4B). Similarly, the timescale of the speed decrease did not depend on the strength of the osmotic shock (Fig. S5C). These results indicate that the loss of motor rotation does not require a specific rotational conformation of the motor. The driving force behind the observed response is independent of the specifics of the stator-rotor interaction, consistent with it being due to a loss of PMF.

**Figure 4.**
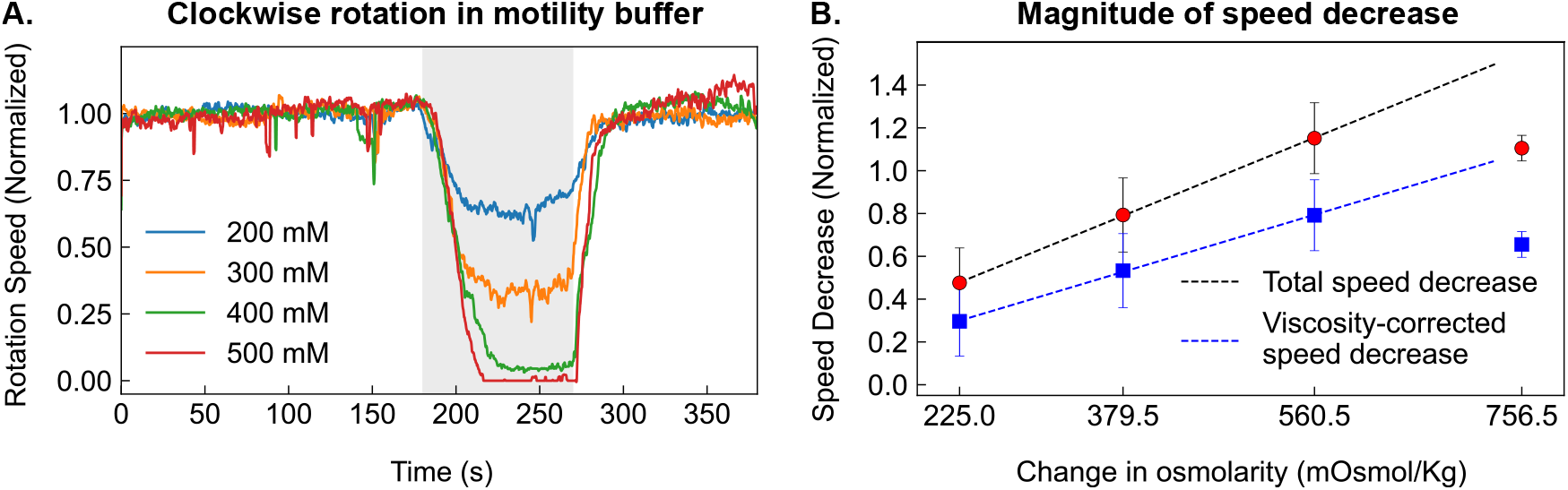
Motor speed reduction is independent of the direction of rotation. **A)** Population-averaged rotation speeds for 200–500 mM sucrose shocks in *E. coli* cells locked in clockwise (CW) rotation (*n*= 8 per condition). Shaded region indicates shock duration. **B)** Normalized magnitude of speed reduction vs. change in osmolarity (from MB with sucrose). The speed goes down to 0 Hz for 500 mM shock, saturating the decrease. Therefore, only values from 200 mM to 400 mM were used to compute the fits. Fit parameters are provided in Table S4.

### TMRM images reveal that a hyperosmotic stress disrupts the membrane potential

While PMF contains contributions both from the membrane potential (Δ*ψ*) and the pH gradient (ΔpH), the former is expected to dominate when the external pH is close to neutral, as in our experiments. Thus, any changes in PMF must be due to changes in membrane potential. To test if the decrease in motor speed during an osmotic shock is associated with depolarization of the membrane, we labeled cells with the Nernstian dye Tetramethylrhodamine methyl ester (TMRM) that reports changes in the membrane potential (Jin et al., 2023; Krasnopeeva et al., 2019). Despite some heterogeneity in labeling and response, we observed many cells in which TMRM intensity decreased during the osmotic shock and recovered after the shock (Fig. S7), with representative examples shown in Fig. 5. The reduced intensity of TMRM during an osmotic shock indicates membrane depolarization. As seen in Fig. 5, the decrease in TMRM intensity was correlated with the strength of the osmotic shock. As negative control, we replaced the media with buffer containing 0 mM additional sucrose, resulting in no change in TMRM intensity. As positive control, we introduced the uncoupler carbonyl cyanide p-trifluoro methoxyphenylhydrazone (FCCP), that resulted in a complete loss of TMRM fluorescence. These data support the idea that an osmotic shock causes disrupts the membrane potential and therefore the PMF, which in turn results in changes in the motor rotation speed.

**Figure 5.**
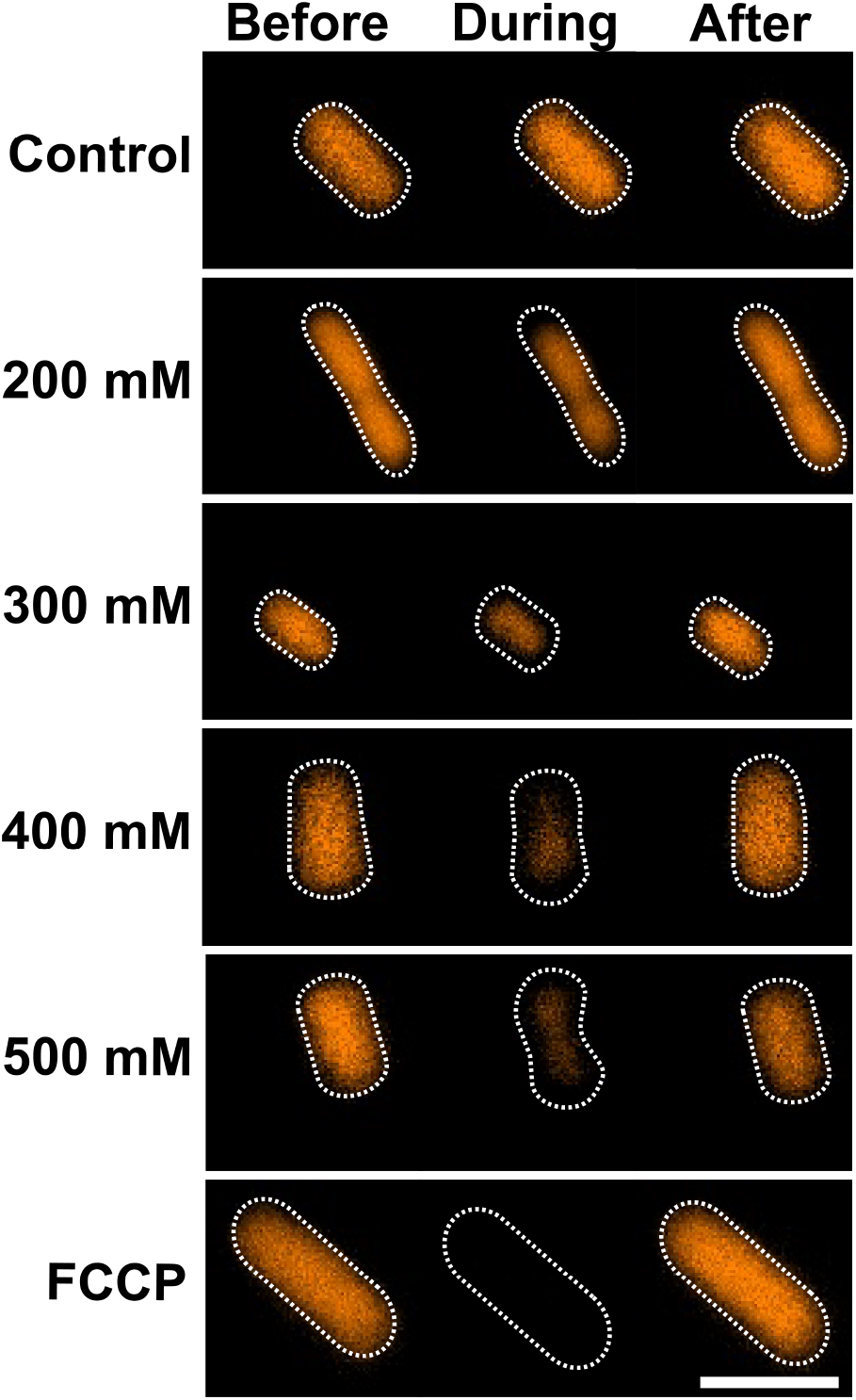
Hyperosmotic shock induces transient membrane depolarization in *E. coli*. Fluorescence images of TMRM-stained cells. Columns show representative cells before, during, and after sucrose shock. **Top row:** Control (0 mM sucrose). **Middle rows:** Cells subjected to 200–500 mM sucrose. TMRM signal drops during the shock and recovers post-shock, with response magnitude increasing with sucrose concentration. **Bottom row:** FCCP-treated cell (100 μM) as a positive control. White dashed outlines show cell boundaries (from phase contrast). Scale bar: 2 *μ*m.

## DISCUSSION

We rationalize that *E. coli*’s response to osmotic shock involves a rapid decrease in PMF. Our bead assay shows that flagellar motor rotation slows immediately upon hyperosmotic stress, with the magnitude of speed loss increasing with shock strength, indicating a dose-dependent response. This effect is independent of the osmolyte used, as both sucrose and sorbitol produced similar results. The response occurred in both counterclockwise and clockwise rotating motors, indicating that the effect originates upstream of the stator-rotor interaction, likely at the level of energy transduction in the stator. Supporting this, TMRM fluorescence decreased during osmotic shock, consistent with PMF loss. We ruled out potassium-induced depolarization, as the response was unchanged in potassium-free media. Together, these results point to PMF loss as a key component of *E. coli*’s response to hyperosmotic stress.

We used the bacterial flagellar motor as a real-time reporter, providing high temporal resolution of PMF. Unlike fluorescence-based methods, which require exogenous dyes that can themselves disrupt PMF (Mancini et al., 2020), motor rotation is native to the cell’s biology and easily accessible from outside. Previous studies have shown that motor speed is linearly proportional to PMF under many conditions (Fung and Berg, 1995; Gabel and Berg, 2003), and PMF is spatially homogeneous within the cell (Biquet-Bisquert et al., 2024), making motor output a reliable readout of the cell’s global energetic state. A recent study found that motor speed plateaus at high PMF and load (Krasnopeeva et al., 2024), indicating a possible non-linear relationship in that regime. However, this does not preclude the motor’s use as a PMF reporter, since any change in motor speed still implies a change in PMF, even if the reverse is not always true under high load.

Leveraging existing knowledge of *E. coli* physiology, we can interpret the time scale of the motor response. The characteristic time extracted from curve fitting (Equation 3) is on the order of a few seconds (Fig. 1F). This is slightly longer than the media exchange time in our flow cells, estimated to be under one second (Berg and Block, 1984). However, osmotic shock induces a rapid decrease in cell volume within seconds (Pilizota and Shaevitz, 2012), suggesting that the motor speed change occurs during this reduction in volume. Whether volume loss directly impacts PMF remains an open question.

Our data suggest that hyperosmotic shock may inhibit stator remodeling. Previous studies have shown that stators disengage when PMF is disrupted (Fung and Berg, 1995; Tipping et al., 2013). In contrast, we observed no evidence of stator remodeling during either the motor slowdown under hyperosmotic stress or its recovery afterward. This is particularly evident in the mismatch between the rapid recovery of motor speed here (a few seconds) and the much slower timescale of stator remodeling (tens of seconds) observed after a period of low load (Wadhwa et al., 2019, 2021). These observations imply that stators remain bound during PMF loss but cease torque generation. One possible explanation is that the rapid volume loss during osmotic shock leads to macromolecular crowding within the cytoplasm (Poolman, 2023), which could reduce stator mobility. However, the interplay between crowding and stator dynamics remains unclear and presents an intriguing direction for future research.

In this study, we examined the impact of hyperosmotic stress, a common and well-characterized bacterial stressor, on the proton motive force (PMF). In *E. coli*, the response to hyperosmotic shock includes the uptake of compatible solutes such as potassium glutamate and glycine betaine (Perroud and Rudulier, 1985). If PMF loss contributes to this response, it raises the question of how PMF-dependent solute transport is maintained. Our results contrast with previous studies that reported no significant changes in PMF, membrane potential, or ATP levels several minutes after osmotic stress was applied (Houssin et al., 1991; Shabala et al., 2009). However, osmotic stress triggers rapid physical and chemical changes within seconds (Pilizota and Shaevitz, 2012), suggesting that a transient PMF drop may have been missed in previous work. The flagellar motor reports PMF with high temporal resolution, revealing fast dynamics that are otherwise overlooked.

Hyperosmotic stress is known to act as a mechanical stimulus, reducing turgor pressure and altering cell shape and size (Pilizota and Shaevitz, 2012, 2013; Rojas, 2020). This triggers the activation of osmoresponsive channels that help restore turgor and cell shape (Sleator and Hill, 2002). Our data suggest that, in addition to these mechanical and structural effects, hyperosmotic stress may also disrupt cellular energetics by causing a rapid loss of PMF. Our findings are in line with those of Rosko et al. (2017), who found that upon encountering an osmotic shock, flagellar rotation rapidly stops, followed by a slower recovery. Our results are also consistent with previous studies showing that mechanical stress can lead to a calcium influx and membrane depolarization (Bruni et al., 2017). However, calcium influx is unlikely to explain our observations, as the media used contained no calcium. While the mechanism of this disruption remains unclear, we speculate that the mechanical strain caused by hyperosmotic stress might disrupt the electron transport chain, leading to the depolarization of the membrane. Elucidating the mechanism behind PMF disruption during osmotic stress represents an exciting avenue for future research.

Together, our results demonstrate that hyperosmotic stress in *E. coli* leads to a rapid, reversible, and dose-dependent loss of PMF, representing an energetic response that adds to well-known mechanical impacts such as loss of turgor and volume reduction. By using the flagellar motor as a high temporal resolution PMF reporter, we uncovered fast energetic dynamics that are likely missed by conventional approaches. These results add a new dimension to our understanding of the bacterial osmotic stress response and open avenues for exploring the interplay between membrane mechanics and energy transduction.

## MATERIALS AND METHODS

### Bacterial Strains, Plasmids, and Cultures

The *E. coli* strain KAF95, a derivative of AW405 was used when testing CCW rotation (Berg and Turner, 1993). Strain KAF95 lacked both the chemotaxis signaling protein CheY and the flagellin protein FliC, and had been transformed with plasmid pFD313 under an ampicillin promoter to express FliC-st. KAF95 cells were grown from a 100 μL frozen stock (stored at *−*80 ^*°*^C) and diluted 1:100 in T-broth (10 g/L tryptone, 5 g/L NaCl) supplemented with 100 μg/mL ampicillin. Cultures were incubated at 33^*°*^C with shaking at 200 rpm until reaching an OD_600_ of 0.5.

For CW rotation, we used strain HCB1797, a derivative of RP437. Briefly, HCB1797 contains an in-frame deletion of FliC in strain VS149 and is transformed with two plasmids: pWB5, which expresses CheY from an isopropyl *β*-D-1-thiogalactopyranoside (IPTG)-inducible promoter, and pKAF131, a chloramphenicol selective plasmid that expresses sticky FliC. HCB1797 cells were grown from a 100 μL frozen stock (stored at *−*80 ^*°*^C) and diluted 1:100 in T-broth (10 g/L tryptone, 5 g/L NaCl) supplemented with 100 μg/mL ampicillin, 25 μg/mL chloramphenicol, and induced with 0.1 mM IPTG. Cultures were incubated at 33^*°*^C with shaking at 200 rpm until reaching an OD_600_ of 0.5.

For fluorescence microscopy assays with TMRM, we used strain MG1655. Cells were grown from a 100 μL frozen stock (stored at *−*80 ^*°*^C) and diluted 1:100 in LB-broth (10 g/L tryptone, 5 g/L NaCl, 5 g/L yeast). Cultures were incubated at 37^*°*^C with shaking at 200 rpm until reaching an OD_600_ of 0.5.

### Bead Assays

Bacterial cultures of either KAF95 or HCB1797 cells were centrifuged at 1,500 *g* for 7 minutes, washed, and resuspended in motility buffer (6.2 mM K_2_HPO_4_, 3.8 mM KH_2_PO_4_, 0.1 mM EDTA, 67 mM NaCl, pH 7.5) or sodium phosphate buffer (6.2 mM Na_2_HPO_4_, 3.8 mM NaH_2_PO_4_, 0.1 mM EDTA, 67 mM NaCl, pH 7.5), supplemented with 60 mM osmolyte to match the osmolarity of the growth medium.

Flagellar filaments were sheared by passing 1 mL of the resuspended culture 150 times through 7 cm of polyethylene tubing (I.D. = 0.58 mm), connected by two syringes equipped with 23-gauge adapters. The cells were then centrifuged again at 1,500 *g* for 7 minutes and resuspended in 400 μL of the baseline buffer. A custom-made flow cell (Berg and Block, 1984) was used to immobilize 15 μL of the cell suspension with sheared flagellar filaments onto a poly-L-lysine-coated coverslip. The flow cell was then inverted for 5 minutes to allow the cells to adhere to the coverslip. The flow chamber was then flushed with 1.0 μm polystyrene beads (0.269% w/v) at a flow rate of 2 mL/min, inverted for 10 minutes to allow bead attachment to the flagellar filaments, and washed again to remove unbound beads.

### Osmotic Shock Assays

Prior to applying the osmotic shock, an initial recording was taken in the baseline media. The shock was applied from 180 s to 270 s by exchanging the baseline media with media supplemented with the desired osmolyte. At 270 s, the media was switched back to the baseline condition, and the recording continued until 400 s. All media exchanges were performed using a Hamilton valve at a constant flow rate of 15 mL/hr for all experiments.

### Fluorescence Microscopy

MG1655 cells grown to an OD_600_ of 0.5 were centrifuged at 1,500 *g* for 7 minutes, washed, and resuspended in a modified version of motility buffer (10 mM K_2_HPO_4_, 10 mM EDTA, 67 mM NaCl, pH 7.5) to permeabilize the outer membrane. The cell suspension was shaken at 200 rpm at room temperature. After 10 minutes, the EDTA was washed out three times by centrifugation at 1,500 *g* for 7 minutes and resuspension in motility buffer. Cells were then diluted 1:10 in MB supplemented with 60 mM sucrose and 0.1 μM TMRM. After an additional 20 minutes of incubation, 15 μL of the cell suspension was added to a poly-L-lysine-coated coverslip mounted in a custom-made flow cell. The flow rate was set to 15 mL/hr to remove any unattached cells prior to imaging. Cells were imaged using a Nikon Eclipse Ti2 microscope equipped with a Plan Apo Lambda 100*×* Oil Ph3 DM objective and a Lumencor Sola II light engine. TMRM fluorescence was visualized using a TRITC filter set with excitation/emission wavelengths of *λ*_ex_ = 540 nm and *λ*_em_ = 579 nm. Images were acquired with a pco.edge camera (Excelitas) at a frame rate of 1.334 s and an exposure time of 10 ms at 10% light intensity. All data acquisition and image capture were performed using the NIS Elements Advanced Research software JOBS module (Nikon Instruments).

### Single Motor Measurments

Raw images of rotating single motors with attached beads were first smoothed using a Gaussian blur to reduce high-frequency noise. The bead region was then segmented using Otsu’s thresholding method, and the resulting mask was converted into binary format for further processing. The centroid of each bead was identified in every frame, and the center of rotation was estimated by fitting a circle to a trajectory of 300 consecutive centroid positions. Rotational frequency was calculated by determining the angular displacement of the bead over time. Finally, the data were normalized by the average pre-shock rotation speed and smoothed with a median filter.

Viscosity values for sucrose and sorbitol solutions were obtained from Swindells et al. (1958) and Sahare (2021), respectively. To estimate the expected change in rotation speed due to changes in viscosity, we used Stokes’ law:

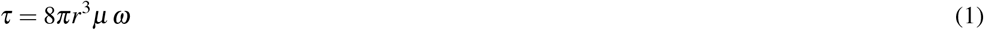

where *τ* is the viscous drag on the bead of radius *r* that is rotating with an angular velocity *ω* in a fluid with viscosity *μ*. If the torque produced by the motor remains constant, then the bead’s rotational speed scales inversely with viscosity, such that

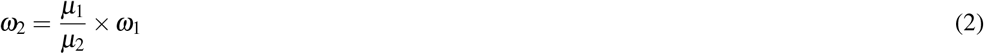

where *ω*_1_ and *ω*_2_ are the angular speeds before and after the viscosity change, and *η*_1_ and *η*_2_ are the corresponding viscosities.

### Curve Fitting

To quantify the magnitude and timescale of the motor speed decrease, we fit the speed of individual motors between 175 and 240 s (Fig. 1C, inset) with an inverse sigmoidal function,

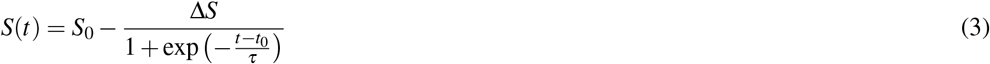

where *S*_0_ is the initial rotation speed, Δ*S* the magnitude of speed decrease, *t* the time in seconds, *t*_0_ the inflection point (half-maximal decrease), and *τ* the characteristic timescale. *S*_0_ and Δ*S* were fixed from average motor speeds measured between 155–175 s, and 215–235 s, respectively. *τ* and *t*_0_ were fit as free parameters.

Each trace was normalized to its mean value during the baseline period preceding the osmotic shock and then fit with Equation 3 to extract the response timescale and magnitude. Fits were performed for individual cells, and the resulting parameters were used to compute population means and standard deviations.

## AUTHOR CONTRIBUTIONS

N.W. designed research; L.M., S.B., and J.Y. performed research; L.M. and E.D. analyzed data; L.M. and N.W. wrote the paper.

## DATA AVAILABILITY

All code and data used in this manuscript are available at the GitHub repository: https://github.com/wadhwalab/hyperosmotic.git.

## ACKNOWLEDGMENTS

We thank Daozheng Gong for preliminary data collection, Shabduli Sawant and Brennen Wise for their assistance with data analysis, and members of the Wadhwa Lab for helpful discussions. We are grateful to Drs. Jeremy Wideman and David Blair for feedback on the manuscript.

## REFERENCES

Akabuogu, E., da Cunha Martorelli, V.C., Roberts, I. S., Waigh, T. A., et al. (2025). Emergence of ion-channel-mediated electrical oscillations in Escherichia coli biofilms. eLife, 13:RP92525.

Altendorf, K., Booth, I. R., Gralla, J., Greie, J.-C., Rosenthal, A. Z., and Wood, J. M. (2009). Osmotic stress. EcoSal Plus, 3(2):10.1128/ecosalplus.5.4.5.

Belas, R. (2014). Biofilms, flagella, and mechanosensing of surfaces by bacteria. Trends in Microbiology, 22(9):517–527.

Benarroch, J. M. and Asally, M. (2020). The microbiologist’s guide to membrane potential dynamics. Trends in Microbiology, 28(4):304–314.

Berg, H. and Turner, L. (1993). Torque generated by the flagellar motor of escherichia coli. Biophysical Journal, 65(5):2201–2216.

Berg, H. C. and Block, S. M. (1984). A miniature flow cell designed for rapid exchange of media under high-power microscope objectives. Microbiology, 130(11):2915–2920.

Biquet-Bisquert, A., Carrio, B., Meyer, N., Fernandes, T. F., Abkarian, M., Seduk, F., Magalon, A., Nord, A. L., and Pedaci, F. (2024). Spatiotemporal dynamics of the proton motive force on single bacterial cells. Science Advances, 10(21):eadl5849.

Biquet-Bisquert, A., Labesse, G., Pedaci, F., and Nord, A. L. (2021). The dynamic ion motive force powering the bacterial flagellar motor. Frontiers in Microbiology, 12:659464.

Bruni, G. N., Weekley, R. A., Dodd, B. J. T., and Kralj, J. M. (2017). Voltage-gated calcium flux mediates Escherichia coli mechanosensation. Proceedings of the National Academy of Sciences, 114(35):9445– 9450.

Buttress, J. A., Halte, M., te Winkel, J. D., Erhardt, M., Popp, P. F., and Strahl, H. (2022). A guide for membrane potential measurements in gram-negative bacteria using voltage-sensitive dyes. Microbiology, 168(9).

Cota-Robles, E. H. (1963). Electron microscopy of plasmolysis in Escherichia coli. Journal of Bacteriology, 85(3):499–503.

Dinnbier, U., Limpinsel, E., Schmid, R., and Bakker, E. P. (1988). Transient accumulation of potassium glutamate and its replacement by trehalose during adaptation of growing cells of Escherichia coli K-12 to elevated sodium chloride concentrations. Archives of microbiology, 150:348–357.

Dufrêne, Y. F. and Persat, A. (2020). Mechanomicrobiology: how bacteria sense and respond to forces. Nature Reviews Microbiology, 18(4):227–240.

Fuchino, K., Flärdh, K., Dyson, P., and Ausmees, N. (2017). Cell-biological studies of osmotic shock response in Streptomyces spp. Journal of Bacteriology, 199(1):e00465–16.

Fung, D. C. and Berg, H. C. (1995). Powering the flagellar motor of Escherichia coli with an external voltage source. Nature, 375(6534):809–812.

Gabel, C. V. and Berg, H. C. (2003). The speed of the flagellar rotary motor of Escherichia coli varies linearly with protonmotive force. Proceedings of the National Academy of Sciences, 100(15):8748–8751.

Gordon, V. D. and Wang, L. (2019). Bacterial mechanosensing: the force will be with you, always. Journal of Cell Science, 132(7):jcs227694.

Houssin, C., Eynard, N., Shechter, E., and Ghazi, A. (1991). Effect of osmotic pressure on membrane energy-linked functions in Escherichia coli. Biochimica et Biophysica Acta (BBA) - Bioenergetics, 1056(1):76–84.

Jin, X., Zhang, X., Ding, X., Tian, T., Tseng, C.-K., Luo, X., Chen, X., Lo, C.-J., Leake, M. C., and Bai, F. (2023). Sensitive bacterial V_m_ sensors revealed the excitability of bacterial V_m_ and its role in antibiotic tolerance. Proceedings of the National Academy of Sciences, 120(3):e2208348120.

Kralj, J. M. et al. (2011). Electrical spiking in Escherichia coli probed with a fluorescent voltage-indicating protein. Science, 333(6040):345–348.

Krasnopeeva, E., Le Nagard, L., Poon, W., Lo, C.-J., and Pilizota, T. (2024). Nonlinear dependency of the bacterial flagellar motor speed on proton motive force and its consequences for swimming. bioRxiv.

Krasnopeeva, E., Lo, C.-J., and Pilizota, T. (2019). Single-cell bacterial electrophysiology reveals mechanisms of stress-induced damage. Biophysical Journal, 116(12):2390–2399.

Lele, P. P., Hosu, B. G., and Berg, H. C. (2013). Dynamics of mechanosensing in the bacterial flagellar motor. Proceedings of the National Academy of Sciences, 110(29):11839–11844.

Lo, C.-J., Leake, M. C., Pilizota, T., and Berry, R. M. (2007). Nonequivalence of membrane voltage and ion-gradient as driving forces for the bacterial flagellar motor at low load. Biophysical Journal, 93(1):294–302.

Maloney, P. C., Kashket, E. R., and Wilson, T. H. (1974). A protonmotive force drives ATP synthesis in bacteria. Proceedings of the National Academy of Sciences, 71(10):3896–3900.

Mancini, L., Terradot, G., Tian, T., Pu, Y., Li, Y., Lo, C.-J., Bai, F., and Pilizota, T. (2020). A general workflow for characterization of Nernstian dyes and their effects on bacterial physiology. Biophysical Journal, 118(1):4–14.

Manson, M. D., Tedesco, P., Berg, H. C., Harold, F. M., and der Drift, C. V. (1977). A protonmotive force drives bacterial flagella. Proceedings of the National Academy of Sciences, 74(7):3060–3064.

Mitchell, P. (1961). Coupling of phosphorylation to electron and hydrogen transfer by a chemiosmotic type of mechanism. Nature, 191:144–148.

Nord, A. L., Gachon, E., Perez-Carrasco, R., Nirody, J. A., Barducci, A., Berry, R. M., and Pedaci, F. (2017). Catch bond drives stator mechanosensitivity in the bacterial flagellar motor. Proceedings of the National Academy of Sciences, 114(49):12952–12957.

Perroud, B. and Rudulier, D. L. (1985). Glycine betaine transport in Escherichia coli: osmotic modulation. Journal of Bacteriology, 161(1):393–401.

Persat, A. (2017). Bacterial mechanotransduction. Current Opinion in Microbiology, 36:1–6.

Pilizota, T. and Shaevitz, J. W. (2012). Fast, multiphase volume adaptation to hyperosmotic shock by Escherichia coli. PLOS ONE, 7(4):1–10.

Pilizota, T. and Shaevitz, J. W. (2013). Plasmolysis and cell shape depend on solute outer-membrane permeability during hyperosmotic shock in E. coli. Biophysical journal, 104(12):2733–2742.

Poolman, B. (2023). Physicochemical homeostasis in bacteria. FEMS Microbiology Reviews, 47(4):fuad033.

Prindle, A., Liu, J., Asally, M., Ly, S., Garcia-Ojalvo, J., and Süel, G.M. (2015). Ion channels enable electrical communication in bacterial communities. Nature, 527(7576):59–63.

Rojas, E., Theriot, J. A., and Huang, K. C. (2014). Response of !i¿escherichia coli!/i¿ growth rate to osmotic shock. Proceedings of the National Academy of Sciences, 111(21):7807–7812.

Rojas, E. R. (2020). The Mechanical Properties of Bacteria and Why they Matter, pages 1–14. Springer International Publishing, Cham.

Rosko, J., Martinez, V. A., Poon, W. C. K., and Pilizota, T. (2017). Osmotaxis in !i¿escherichia coli!/i¿ through changes in motor speed. Proceedings of the National Academy of Sciences, 114(38):E7969– E7976.

Sahare, S. P. (2021). Studies of molecular interactions of sugar alcohols in water by volumetric and visco-metric measurement. International Journal of Scientific Research in Science and Technology(IJSRST), 8(1):422–424.

Shabala, L., Bowman, J., Brown, J., Ross, T., McMeekin, T., and Shabala, S. (2009). Ion transport and osmotic adjustment in Escherichia coli in response to ionic and non-ionic osmotica. Environmental Microbiology, 11(1):137–148.

Sleator, R. D. and Hill, C. (2002). Bacterial osmoadaptation: the role of osmolytes in bacterial stress and virulence. FEMS Microbiology Reviews, 26(1):49–71.

Stautz, J., Hellmich, Y., Fuss, M. F., Silberberg, J. M., Devlin, J. R., Stockbridge, R. B., and Hänelt, I. (2021). Molecular mechanisms for bacterial potassium homeostasis. Journal of Molecular Biology, 433(16):166968.

Stephens, C., Martinez, M., Leonardi, V., Jaing, J., and Miller, A. (2024). The scr and csc pathways for sucrose utilization co-exist in E. coli, but only the scr pathway is widespread in other enterobacteriaceae. Frontiers in Microbiology, 15:1409295.

Strahl, H. and Hamoen, L. W. (2010). Membrane potential is important for bacterial cell division. Proceedings of the National Academy of Sciences, 107(27):12281–12286.

Swindells, J. F., Snyder, C., Hardy, R. C., and Golden, P. (1958). Viscosities of sucrose solutions at various temperatures: Tables of recalculated values, volume 440. US Government Printing Office.

Tipping, M. J., Delalez, N. J., Lim, R., Berry, R. M., and Armitage, J. P. (2013). Load-dependent assembly of the bacterial flagellar motor. mBio, 4(4):10.1128/mbio.00551-13.

Wadhwa, N. and Berg, H. C. (2022). Bacterial motility: machinery and mechanisms. Nature reviews microbiology, 20(3):161–173.

Wadhwa, N., Phillips, R., and Berg, H. C. (2019). Torque-dependent remodeling of the bacterial flagellar motor. Proceedings of the National Academy of Sciences, 116(24):11764–11769.

Wadhwa, N., Tu, Y., and Berg, H. C. (2021). Mechanosensitive remodeling of the bacterial flagellar motor is independent of direction of rotation. Proceedings of the National Academy of Sciences, 118(15):e2024608118.

Wong, F. and Amir, A. (2019). Mechanics and dynamics of bacterial cell lysis. Biophysical Journal, 116(12):2378–2389.

Yuan, J., Fahrner, K. A., Turner, L., and Berg, H. C. (2010). Asymmetry in the clockwise and counter-clockwise rotation of the bacterial flagellar motor. Proceedings of the National Academy of Sciences, 107(29):12846–12849.

