## SUPPLEMENTARY MATERIAL for "Osmotic stress triggers fast and reversible PMF collapse in *Escherichia coli*"

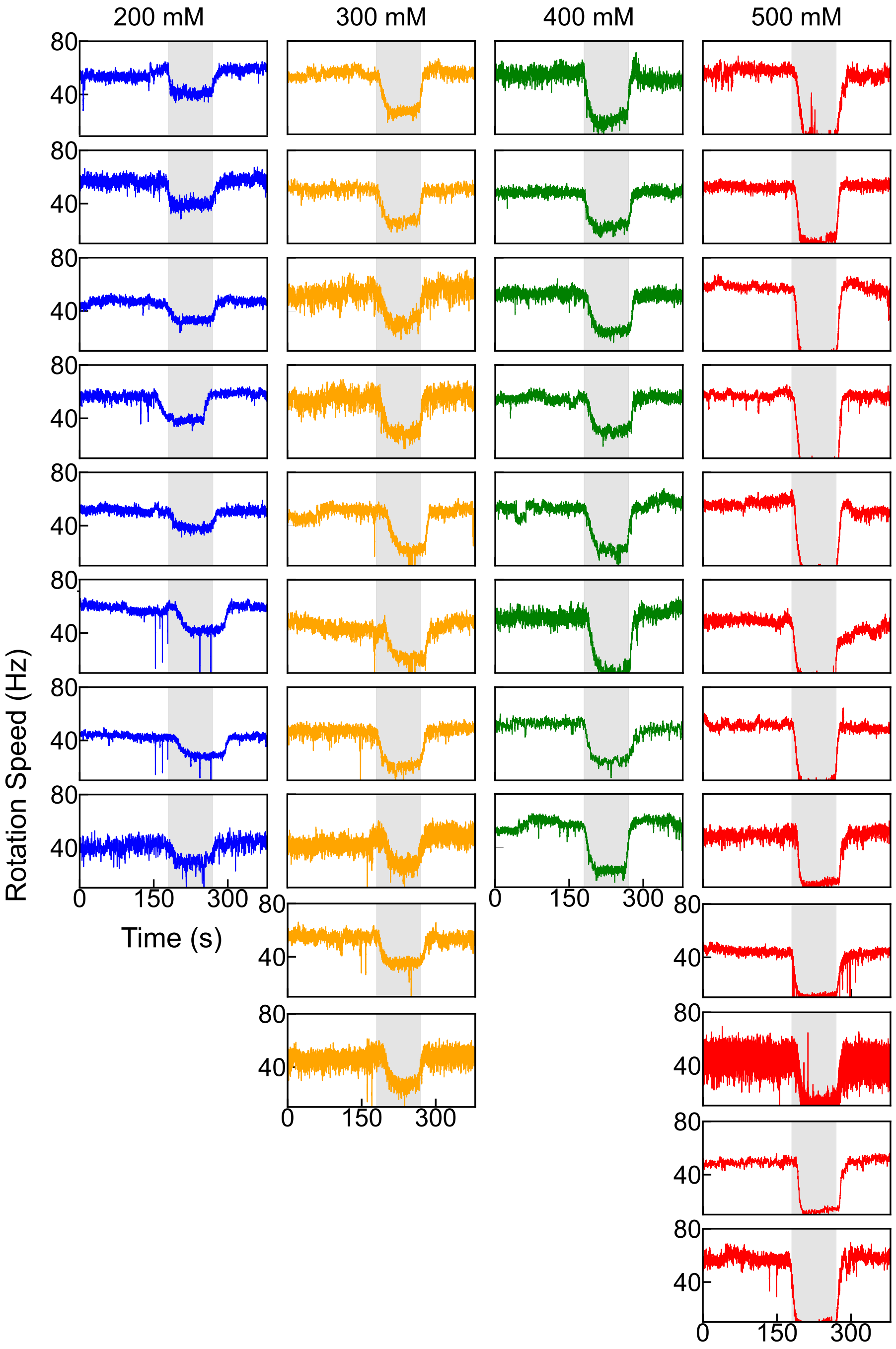


**Supplementary Figure S1. Individual motor speed traces from the sucrose osmotic shocks.** The four columns correspond to sucrose shocks of 200 mM, 300 mM, 400 mM, and 500 mM. The shaded region represents the duration of the shock.

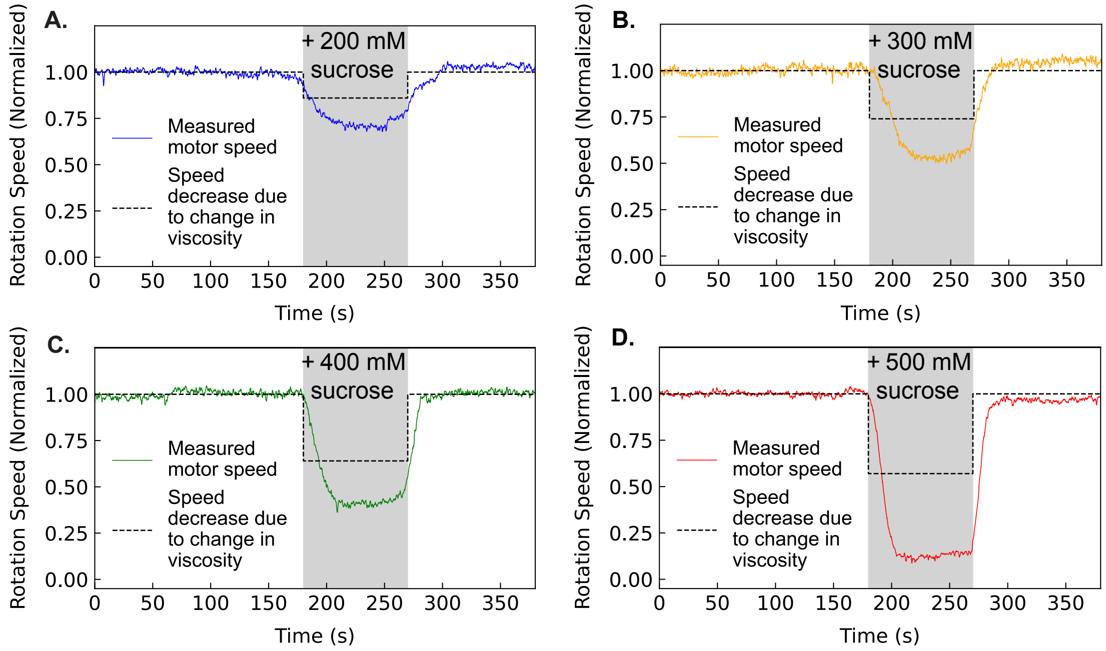


**Supplementary Figure S2. Motor speed reduction during osmotic shock exceeds that expected from increased viscosity alone. A–D)** Population-averaged normalized rotation speeds during 200, 300, 400, and 500 mM sucrose shocks ($n$ = 8, 10, 8, and 12, respectively). Shaded regions indicate the duration of hyperosmotic exposure. Dashed black lines show the estimated changes in speed due solely to increased viscosity, calculated using Stokes' law. Estimated normalized speed reductions for 200, 300, 400, and 500 mM sucrose are 0.18, 0.26, 0.36, and 0.43, respectively. Observed speed reductions exceed these values, indicating a contribution from PMF dissipation beyond viscosity effects.

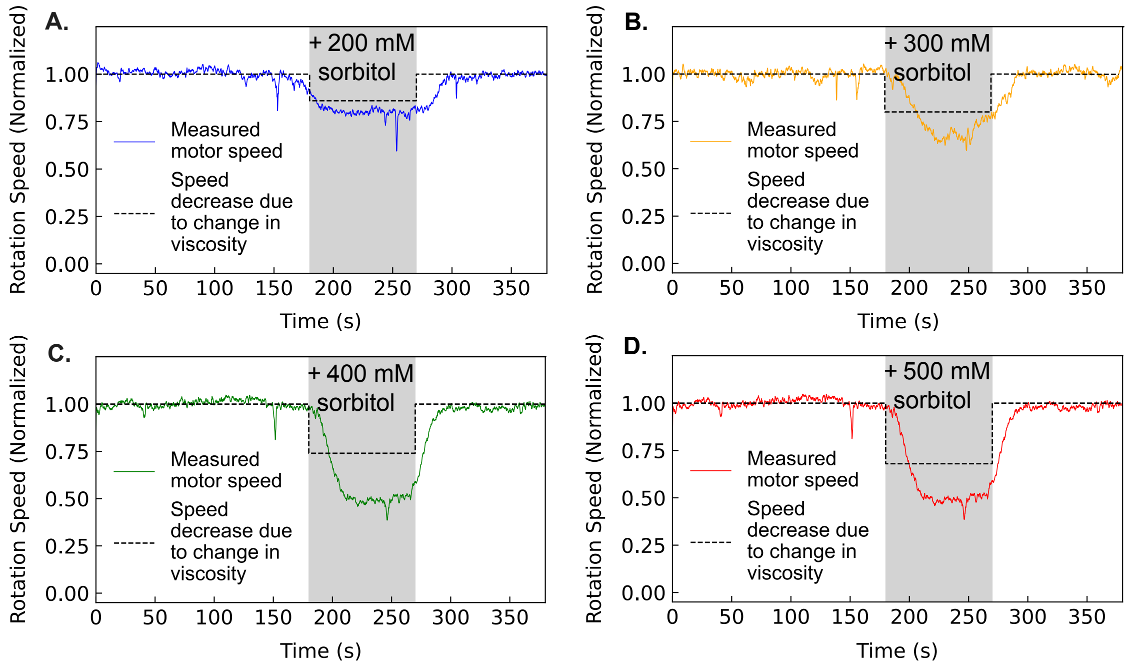


**Supplementary Figure S3. Motor speed reduction during osmotic shock exceeds that expected from increased viscosity alone. A–D)** Population-averaged normalized rotation speeds during 200, 300, 400, and 500 mM sorbitol shocks ($n$ = 8). Shaded regions indicate the duration of hyperosmotic exposure. Dashed black lines show the estimated changes in speed due solely to increased viscosity, calculated using Stokes' law. Estimated normalized speed reductions for 200, 300, 400, and 500 mM sorbitol are 0.14, 0.20, 0.26, and 0.32, respectively. Observed speed reductions exceed these values, indicating a contribution from PMF dissipation beyond viscosity effects.

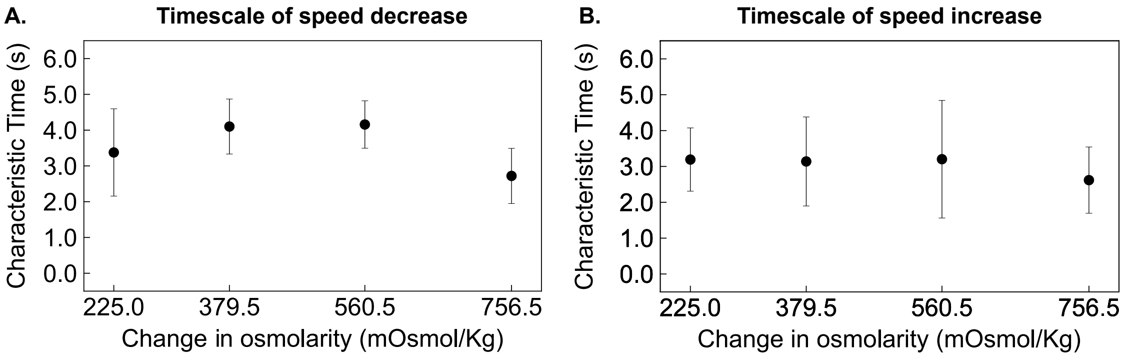


**Supplementary Figure S4. Timescale of rotation speed as a function of change in osmolarity (from sucrose in MB) A)** Characteristic time for speed decrease during the decline phase, with black circles and error bars indicating mean and SD, respectively. **B)** Characteristic time for speed increase during the recovery phase, with black circles and error bars indicating mean and SD, respectively. Characteristic time shows no significant dependence on shock for either the decline or recovery phase.

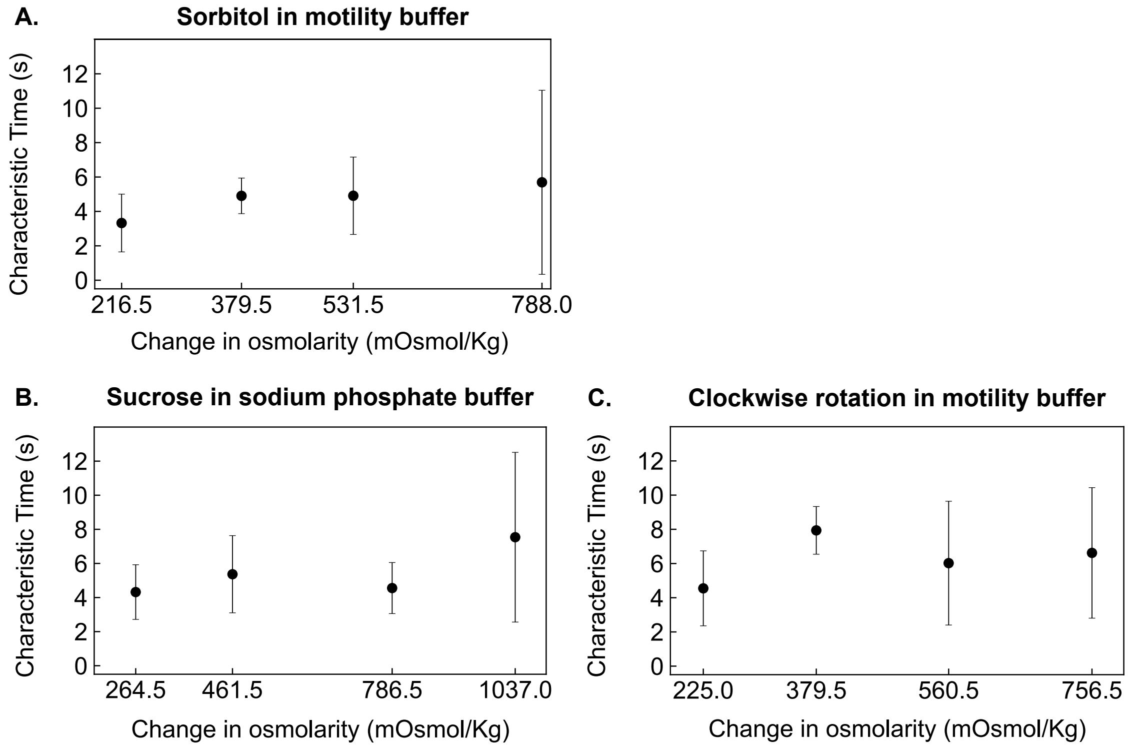


**Supplementary Figure S5. Timescale of speed decrease as a function of change in osmolarity.** **A–C)** Characteristic time representing the timescale of speed decrease for: (**A**) sorbitol in MB, (**B**) sucrose in SPB, and (**C**) clockwise rotation of sucrose in MB. Black circles and error bars indicate the mean and SD, respectively. Characteristic time shows no significant dependence on shock strength.

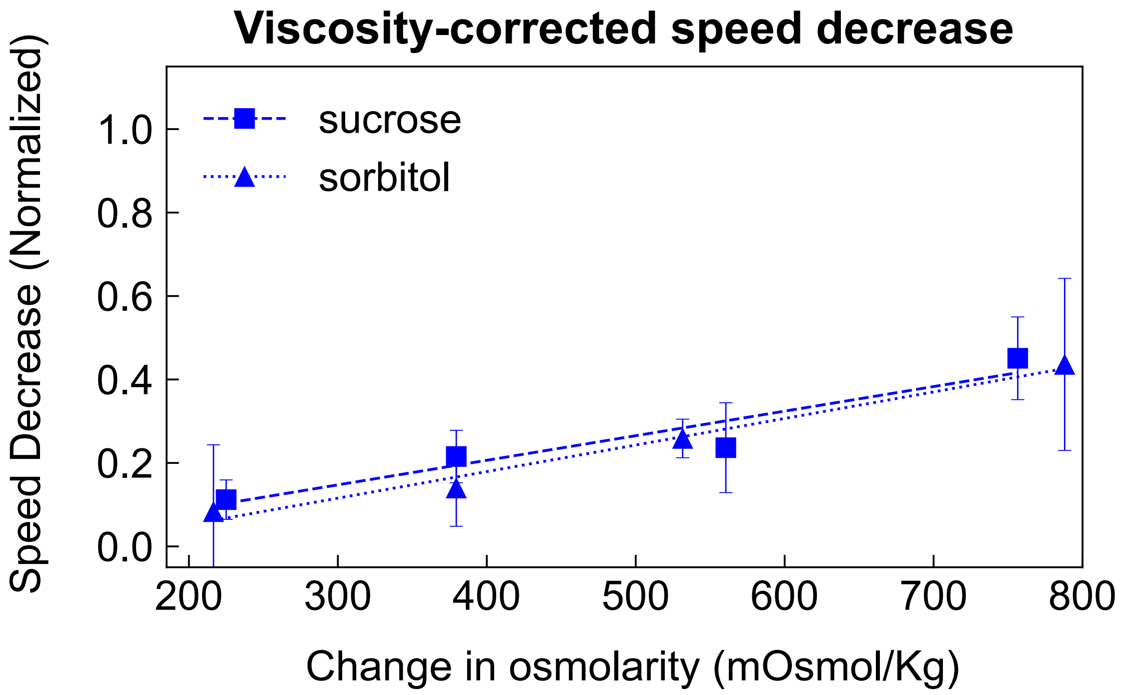


**Supplementary Figure S6. Normalized speed reduction versus change in osmolarity for sucrose (dashed line) and sorbitol (dotted line).** Blue markers and error bars represent the difference between the total speed decrease and the estimated reduction due to changes in viscosity. The dashed and dotted lines show linear fits to this difference. Statistical values for the linear fits are provided in (Table *S4*).

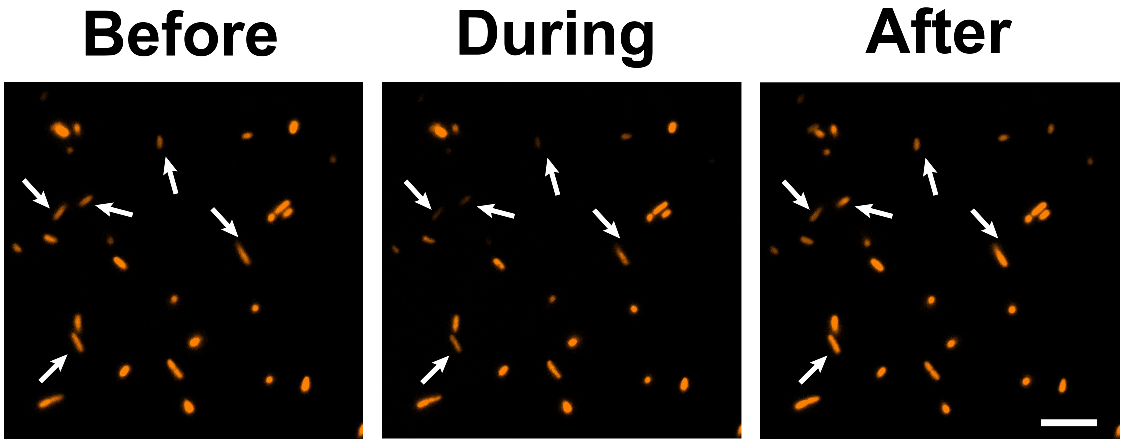


**Supplementary Figure S7. Fluorescence images of heterogeneously labeled TMRM-stained cells.** Columns show representative cells before, during, and after a 300 mM sucrose shock. White arrows point to cells that follow the decrease in fluorescence during the shock and recover afterwards. Scale bar: 10 μm.

| Sucrose Concentration (mM) | Viscosity (cP) |
|---|---|
| 60 | 1.053 |
| 260 | 1.291 |
| 360 | 1.423 |
| 460 | 1.650 |
| 560 | 1.861 |

**Supplementary Table S1. Viscosity of MB supplemented with sucrose.** Viscosity values obtained from Swindells et al. (1958)..

| Sorbitol Concentration (mM) | Viscosity (cP) |
|---|---|
| 60 | 1.053 |
| 260 | 1.227 |
| 360 | 1.323 |
| 460 | 1.419 |
| 560 | 1.540 |

**Supplementary Table S2. Viscosity of MB supplemented with sorbitol.** Viscosity values obtained from Sahare (2021).

| Condition | $m$ ($\times 10^{-4}$) | $b$ | $R^2$ |
|---|---|---|---|
| Sucrose | 4.7 | 0.078 | 0.99 |
| Sorbitol | 3.1 | 0.079 | 0.98 |

**Supplementary Table S3. Linear fit parameters for the estimated speed decrease due to changes in viscosity of sucrose and sorbitol solutions.**

| | Total speed decrease | | | Corrected speed decrease | | |
|---|---|---|---|---|---|---|
| Condition | $m$ ($\times 10^{-3}$) | $b$ | $R^2$ | $m$ ($\times 10^{-3}$) | $b$ | $R^2$ |
| Sucrose in MB | 1.1 | 0.049 | 0.98 | 0.48 | 0.078 | 0.99 |
| Sorbitol in MB | 0.95 | 0.0032 | 0.99 | 0.64 | −0.075 | 0.98 |
| Sucrose in SPB | 0.69 | 0.25 | 0.97 | 0.34 | 0.16 | 0.91 |
| Clockwise rotation MB | 2.0 | 0.025 | 1.0 | 1.5 | −0.033 | 1.0 |

**Supplementary Table S4. Linear fit parameters for each experimental condition.** The total speed decrease was determined from the mean and SD for each shock concentration. The corrected speed decrease was obtained by subtracting the estimated reduction in speed due to viscosity from the total speed decrease.

| Concentration (mM) | Change in osmolarity (mOsmol/kg) | | |
|---|---|---|---|
| | Sucrose in MB | Sorbitol in MB | Sucrose in SPB |
| 60 | 0 | 0 | 0 |
| 260 | 225 | 227 | 265 |
| 360 | 380 | 380 | 462 |
| 460 | 561 | 532 | 787 |
| 560 | 757 | 788 | 1040 |

**Supplementary Table S5. Changes in osmolarity for various concentration of sucrose and sorbitol in MB and SPB.** Osmolarity measurements were performed using a vapor pressure osmometer (OSMETTE II™, Precision Systems Inc., Natick, MA, USA).
